# Isolation and Quantification of mRNAs from Subcellular Phase-Separated Structures from Detergent Permeabilized Brain Cells

**DOI:** 10.1101/2025.08.06.669016

**Authors:** Sritama Ray, Kamalika Mukherjee, Suvendra N. Bhattacharyya

## Abstract

Post-transcriptional regulation by RNA processing bodies, also known as P-bodies (PBs), is crucial for mRNA translation, localization, and stability in various animal cells, including neurons and glial cells. PBs facilitate spatial regulation of protein synthesis, influencing differentiation and synaptic function. Understanding mRNA dynamics in PBs is vital and is suspected to be impaired in neurodegenerative disorders. We developed a detergent-based method to isolate phase-separated P-bodies (PBs) free from cytosolic factors and RNAs, which allows us to examine their mRNA content under altered conditions. Using neuronal and glial cell models, we investigated the impact of Aβ-oligomers on PB-associated mRNAs. In neurons, exposure to amyloid proteins disrupted the release of neuronal differentiation-related mRNAs, hindering their translation. Glial cells exhibited increased levels of cytoplasmic pro-inflammatory cytokine mRNAs after amyloid treatment, as they escaped from RNA processing bodies (PBs), ensuring enhanced translation. These findings underscore the dual role of PBs in controlling mRNA dynamics across cell types and illustrate how amyloid-induced stress may differently affect PB-mediated post-transcriptional control in neurons and astroglia. This method provides a platform for studying the mechanism and quantifying sub-organellar mRNA localization, as well as its effects on altered gene expression caused by amyloid proteins.

**Graphical Abstract:** 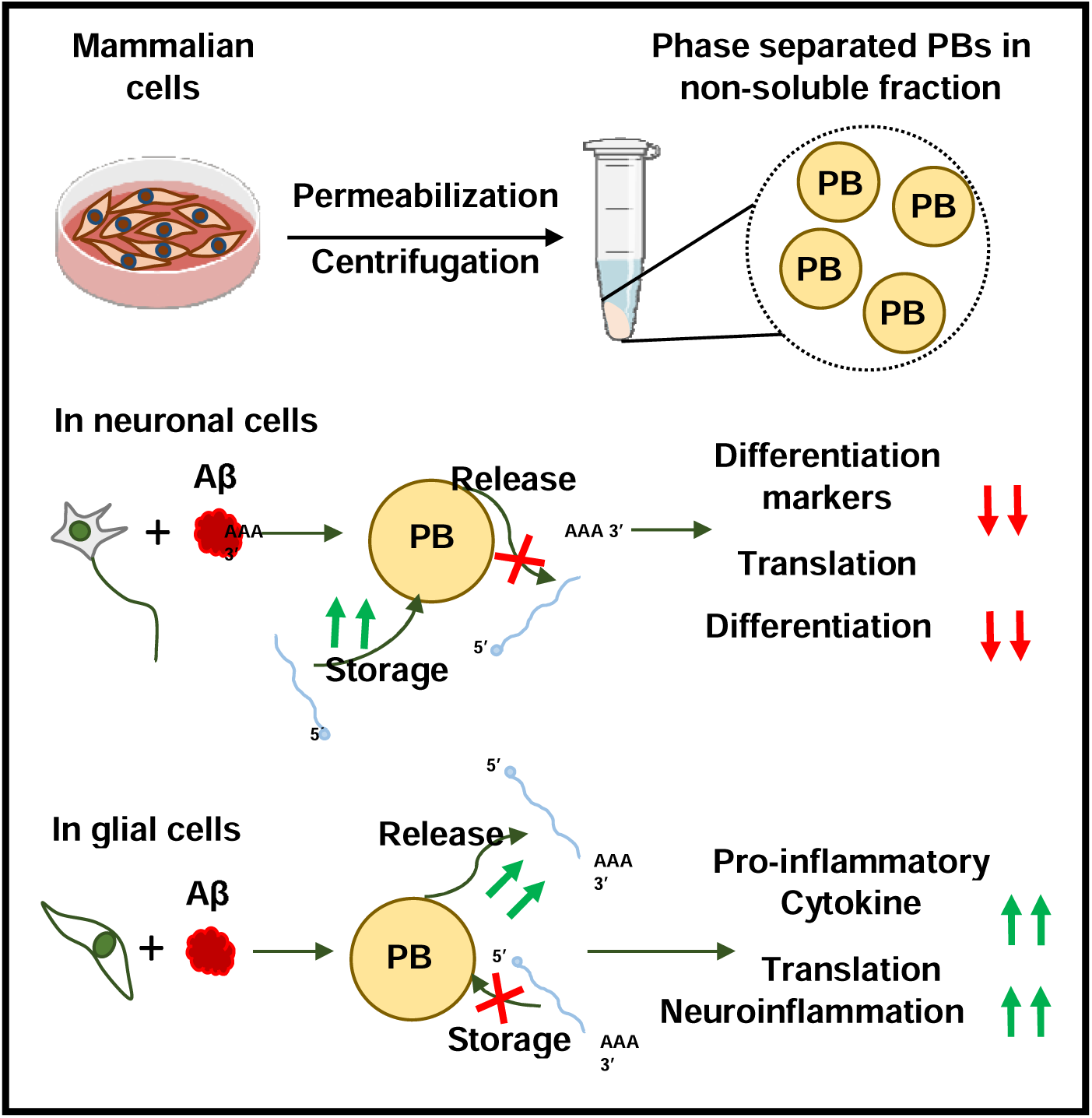

**Highlights:**

- A detergent-based differential solubilization of RNAs from PBs in brain cells.
- In neuronal cells, β-Amyloid oligomers target specific neuronal differentiation-related mRNAs to PBs.
- In astroglia, β-amyloid triggers the escape of inflammatory cytokine mRNAs from PBs.
- Altered mRNAs -protein phase separation connects PBs to neuroinflammation.

## Introduction

RNA regulation at pre- and post-transcriptional levels is influenced by multiple aspects and regulatory factors that orchestrate protein expression at specific times and in particular compartments in metazoan cells. This spatiotemporal control of protein synthesis is governed by the accessibility of mRNAs encoding the protein for translational machineries, which is regulated by various RNA regulatory factors that bind to specific sequences or motifs of the mRNAs through direct or indirect (miRNA-mediated) interactions (Filipowicz et al., 2008; Gebauer and Hentze, 2004). These interactions control translatability, storage, and degradation of mRNAs. RNA-protein interactions are often linked to the formation of phase-separated structures involving complex biophysical and biochemical interactions that result in phase-separated mRNAs, which exhibit reduced accessibility for translating polysomes and the translational machinery(Lin et al., 2015; Van Treeck and Parker, 2018). This process facilitates mRNA storage for specific cellular needs or degradation after repression by miRNAs in diverse mammalian cell types, including neurons(Cougot et al., 2008; Sephton and Yu, 2015).

Among the many regulatory factors involved, microRNAs (miRNAs) play a central role. These small RNA molecules regulate gene expression by binding to complementary regions of target messenger RNAs (mRNAs)(Bartel, 2009). This interaction can either promote the degradation of mRNA or render it translationally inactive, depending on the degree of complementarity between the miRNA and its target sequence. Perfect binding activates the Argonaute protein Ago2, a component of the miRISC complex, which mediates target mRNA degradation through its endonuclease activity. In contrast, imperfect binding leads to the storage of target mRNA in a translationally inactive state within specialized phase-separated structures in the cell(Bartel, 2018; Filipowicz et al., 2008; Jo et al., 2015; Jonas and Izaurralde, 2015).

A critical element of this regulation is RNA processing bodies (PBs), also known as P-bodies. These cytoplasmic, membrane-less RNA-protein condensates are formed through a process called liquid-liquid phase separation(Parker and Sheth, 2007). PBs function as storage hubs for mRNAs bound by miRNA-induced silencing complexes (miRISC), leading to the translational repression of these mRNAs(Liu et al., 2005; Pillai et al., 2004). Importantly, PBs are dynamic structures rather than static ones, with mRNAs constantly shuttling between these bodies, where mRNAs are stored in an inactive form, and polysomes, where mRNAs are actively translated into proteins. The movement of mRNAs between these two compartments is sensitive to changes in the cellular environment, such as those caused by cellular stress or other extrinsic and intrinsic signals, which can directly impact the regulation of mRNA translation and the cellular protein synthesis machinery (Bhattacharyya et al., 2006).

This dynamic regulation of mRNA translation and localization is particularly critical for neurons. Neurons with specialized morphological structures need the regulation of protein synthesis at specific locations within the cell. The localized translation of proteins in particular regions of the neuron, such as dendrites or axonal terminals, is crucial for maintaining synaptic function and plasticity (Holt et al., 2019). In neurons, many proteins required for synaptic function are not synthesized in the cell body, where they remain translationally repressed within PBs. These mRNAs are then transported to distal dendritic regions, where they are released from repression and translated into the proteins necessary for synaptic activities (Cougot et al., 2008). This spatiotemporal control of mRNA translation prevents unnecessary (and potentially harmful) accumulation of proteins in areas where they are not needed, thereby optimizing neuronal function and maintaining cellular homeostasis.

However, the processes regulating the localization of mRNAs to PBs or translating polysomes are responsive to environmental cues, including neuronal activity and stress factors (Cougot et al., 2008; Filipowicz et al., 2008). Environmental changes can impact cellular homeostasis, and disruptions in these regulatory processes may contribute to the development of disease. Despite substantial progress in understanding mRNA localization and translation regulation, the mechanisms driving the differential localization of mRNAs to PBs and the factors influencing this process have remained active research areas over the last two decades. In this context, our study aimed to develop an assay system that employs detergent-based separation of cytosolic content to isolate the insoluble fraction containing phase-separated P-bodies (PBs), thus enabling the analysis of their mRNA content in various cell types within different cellular environments. This will be particularly valuable for studying mRNAs involved in neuronal differentiation and for identifying factors that influence their localization to P-bodies (PBs).

Amyloid proteins and peptides like Aβ_1-42_ form oligomers that cause neurodegeneration(Butterfield and Boyd-Kimball, 2004). These amyloid oligomers, linked to neurodegenerative diseases like Alzheimer’s Disease (AD), also disrupt miRNA function, impairing gene expression regulation. This dysfunction leads to the expression of pro-inflammatory cytokines in astroglial cells, contributing to the inflammation observed in AD (De and Bhattacharyya, 2021). How do these structures regulate mRNA translation into proteins in astroglial cells or neurons? Our assays showed that amyloid oligomers ensure the retention of pro-inflammatory cytokine mRNAs, such as IL-6 and IL-1β, with polysomes by keeping them out of processing bodies (PBs)—likely by inhibiting their interaction with repressing miRNA-Ago protein complexes (miRNPs), which remain associated with PBs after Aβ treatment, leading to target cytokine mRNAs accumulation with polysomes and promotes inflammatory response observed in neurodegenerative diseases(De and Bhattacharyya, 2021; De et al., 2021). Notably, in NGF-differentiated rat pheochromocytoma PC12 cells, a model for studying Aβ_1-42_’s effects on sympathetic neurons(Fujita et al., 1989; Mullenbrock et al., 2011), the results differ. We found that with NGF-induced differentiation, mRNAs vital for neuronal differentiation, like GAP43, Neurofilament-M, and HuD, are targeted to PBs in differentiated PC12 cells exposed to Aβ_1-42_. In the absence of Aβ_1-42_ these mRNAs are released from PBs and translated into proteins necessary for synaptic function during differentiation. Thus, Aβ_1-42_ oligomers promote PB targeting of mRNAs, impairing neuronal differentiation.

This study highlights the intricate relationship between mRNA localization to P-bodies with post-transcriptional regulation in the context of neuronal differentiation and neuroinflammatory responses caused by pathogenic proteins. The findings highlight the role of amyloid oligomers in disrupting the regulatory processes of mRNA targeting to P-bodies (PBs), providing insight into a deeper understanding of how the dysregulation of RNA processing and translation contributes to disease pathology. Our research offers new insights into potential therapeutic strategies targeting RNA regulation, which could have significant implications for diseases characterized by disrupted RNA homeostasis, such as neurodegenerative disorders (e.g., Alzheimer’s disease) or chronic inflammatory conditions. Future research in this area will further elucidate the mechanisms governing mRNA localization and translation, offering potential pathways for developing novel treatments for these challenging diseases. The method described here for studying PB-associated mRNAs is qualitatively and, more importantly, quantitatively essential to reveal the mechanism of phase-separation control of mRNAs in neuronal and non-neuronal cells.

## Results

### Enrichment of phase-separated structures in detergent-permeabilized cells depleted of cytosolic content

In this study, we aimed to develop a biochemical method using digitonin-mediated selective permeabilization of the plasma membrane to remove cytosolic content through leaks created by the detergent. This was followed by the specific isolation of phase-separated, insoluble Dcp1a-positive RNA processing bodies (PBs) from control and NGF-differentiated PC12 cells. This approach enabled us to systematically examine changes in the association of target mRNAs with Dcp1a bodies during neuronal differentiation by removing signals from free cytosolic mRNAs (**Fig. 1A**). The experiment involved selectively permeabilizing the cell membrane with digitonin, which allowed the release of cytosolic content without significantly disrupting cellular structures and organelles. We previously used this method to produce ghost cells with intact P-bodies, mitochondria, and ER structures while also allowing the free passage of externally added molecules to study protein dynamics in the context of phase separation and PB association (Ray et al., 2025). Increasing digitonin concentrations start to affect the integrity and number of PBs in both non-differentiated and differentiated PC12 cells (**Fig. S1A, S1B, and Fig. 1B, C**). After assessing β-tubulin release levels in digitonin-treated cells, we chose 50 ng/mL as the optimal condition because it maintains PB integrity while allowing cytosolic protein leakage to enrich organellar structures and PBs in detergent-treated cells. Western blot analyses showed a gradual decrease in soluble cytosolic proteins with higher detergent concentrations in the insoluble pellet (**Fig. 1D, E**). Through optimization, we found that 50 ng/mL digitonin effectively permeabilizes both undifferentiated and differentiated PC12 cells, extracting a significant portion of soluble proteins while preserving the integrity of the non-soluble, plasma membrane-associated fraction, which is enriched in proteins and mRNA. This fraction provides a starting material for PB-associated RNA extraction.

**Figure 1.**
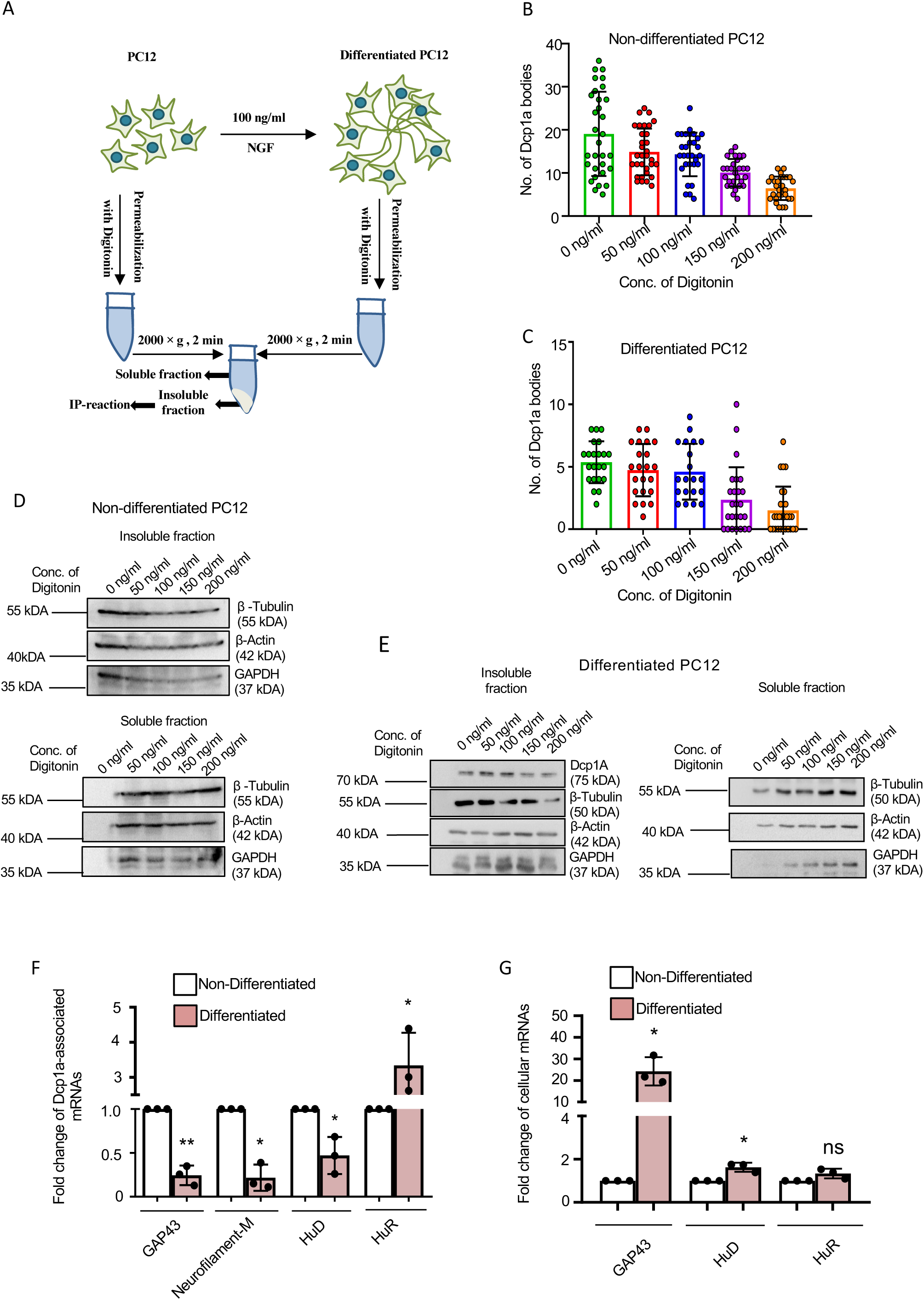
Detergent permeabilization of PC12 cells to enrich RNA processing bodies for downstream extraction and analysis. **(A)** Schematic representation showing separation of soluble and non-soluble fractions from undifferentiated or 72-hour differentiated PC12 cells permeabilized with Digitonin. **(B)** Quantification of GFP-Dcp1a bodies in undifferentiated PC12 cells permeabilized with increasing concentration of Digitonin (0, 50, 100, 150, 200 ng/mL) for 10 minutes at 4 °C. **(C)** Quantification of GFP-Dcp1a bodies in 72-hour differentiated (with 100 ng/mL NGF) PC12 cells permeabilized with increasing concentration of Digitonin (0, 50, 100, 150, 200 ng/mL) for 10 minutes at 4 °C. **(D)** Western blots of endogenous Dcp1a, β-Tubulin, β-Actin, and GAPDH were done to measure the expression levels of these proteins in the non-soluble fraction (upper panel) and soluble fraction (lower panel) of undifferentiated PC12 cells permeabilized with increasing concentration of Digitonin (0, 50, 100, 150, 200 ng/mL). Molecular weight markers are shown. **(E)** Western blot of endogenous Dcp1a, β-Tubulin, β-Actin and GAPDH were done to measure the expression levels of these proteins in the non-soluble fraction (upper panel) and soluble fraction (lower panel) of 72 hours differentiated (with 100 ng/mL NGF) PC12 cells permeabilized with increasing concentration of Digitonin (0, 50, 100, 150, 200 ng/mL). Molecular weight markers are shown. **(F)** qRT-PCR-based quantification of associated GAP43, Neurofilament-M, HuD, and HuR mRNA levels normalized against relative pulled-down Dcp1a levels. Immunoprecipitation of endogenous Dcp1a protein was performed from undifferentiated or 72-hour NGF-differentiated PC12 cells that were permeabilized with 50 ng/ml Digitonin. [p=0.0071 (for GAP43), p=0.0120 (for Neurofilament-M), p=0.0495 (for HuD), p=0.0490 (for HuR), n=3 independent experiments]. **(G)** qRT-PCR based quantification of cellular GAP43, HuD, and HuR mRNA levels normalized against GAPDH level in total RNA isolated from undifferentiated or 72-hour NGF-differentiated PC12 cells [p=0.0254 (for GAP43), p=0.0352 (for HuD), p=0.1195 (for HuR), n=3 independent experiments]. Data represents means ± SDs; ns, non-significant, *p < 0.05, **p< 0.01, ***p < 0.001, ****p < 0.0001. p-values were obtained by using a two-tailed paired Student’s t-test.

### Isolation and quantification of RNA associated with phase-separated components from naïve and differentiated PC12 cells

To explore the differential association of specific mRNAs encoding neuronal differentiation markers and RNA-binding proteins (RBPs) with PBs, we performed immunoprecipitation of the PB-associated protein Dcp1a from both undifferentiated and differentiated PC12 cells. Dcp1a is a PB marker, and solubilization of cytoplasmic content by digitonin should enrich for PB-localized Dcp1a-associated mRNAs in the immunoprecipitation reaction performed with Dcp1a-specific antibodies (**Fig. 1A; F-G**). We focused on four target mRNAs: two encoding key differentiation markers—GAP43 and Neurofilament-M—which are expected to increase in expression in differentiated PC12 cells, and two encoding RBPs—the neuron-specific HuD and the ubiquitously expressed HuR. Our analysis of cellular and Dcp1a-associated mRNAs indicated enrichment of HuR mRNAs associated with Dcp1a, while the association of GAP43, HuD, and Neurofilament-M mRNAs with Dcp1a decreased, despite their increased expression during differentiation. This supports the hypothesis that PB targeting of mRNA and its interaction with PB components, such as Dcp1a, are inversely regulated. Quantitative RT-PCR showed that during neuronal differentiation, mRNAs for differentiation markers such as GAP43 and Neurofilament-M were significantly released from PBs, reflecting derepression and active translation supporting differentiation. Similarly, the association of mRNA encoding neuron-specific RBP HuD, with PBs decreased during differentiation, suggesting increased availability for translation. Conversely, the mRNA for HuR, a ubiquitously expressed RBP involved in cell growth and proliferation, showed increased localization to PBs after differentiation, likely reflecting the growth arrest necessary for proper neuronal differentiation (**Fig. 1F and G**).

### Amyloid beta oligomers affect mRNA association with phase-separated PB structures in differentiated PC12 cells

To investigate how amyloid proteins interfere with the neuronal differentiation process, we treated differentiated PC12 cells with 0 and 2.5 µM amyloid beta Aβ(1-42) and then immunoprecipitated Dcp1a from digitonin-permeabilized cells (**Fig. 2A**). Differentiated PC12 cells treated with digitonin also retain Ago2 and RCK/p54 in the Dcp1a-positive bodies used for Dcp1a IP and RNA extraction (**Fig. 2B and C**). Dcp1a was found to be associated with RCK/p64, a classic marker protein of P-bodies (PBs), in immunoprecipitation (**Fig. 2B**). RNA analysis revealed a significant increase in the association of GAP 43, Neurofilament-M, and HuD mRNAs-transcripts that were found to dissociate from PBs during NGF-mediated PC12 differentiation (**Fig. 1F-G**)-with PBs isolated following Aβ(1-42) treatment (**Fig. 2D and E**). These results suggest that Aβ(1-42) disrupts the standard regulatory mechanisms controlling mRNA escape from PBs in differentiating PC12 cells and promotes the translational repression of differentiation-related protein-coding mRNAs such as Neurofilament-M, GAP 43, and HuD in cells treated with amyloid protein.

**Figure 2.**
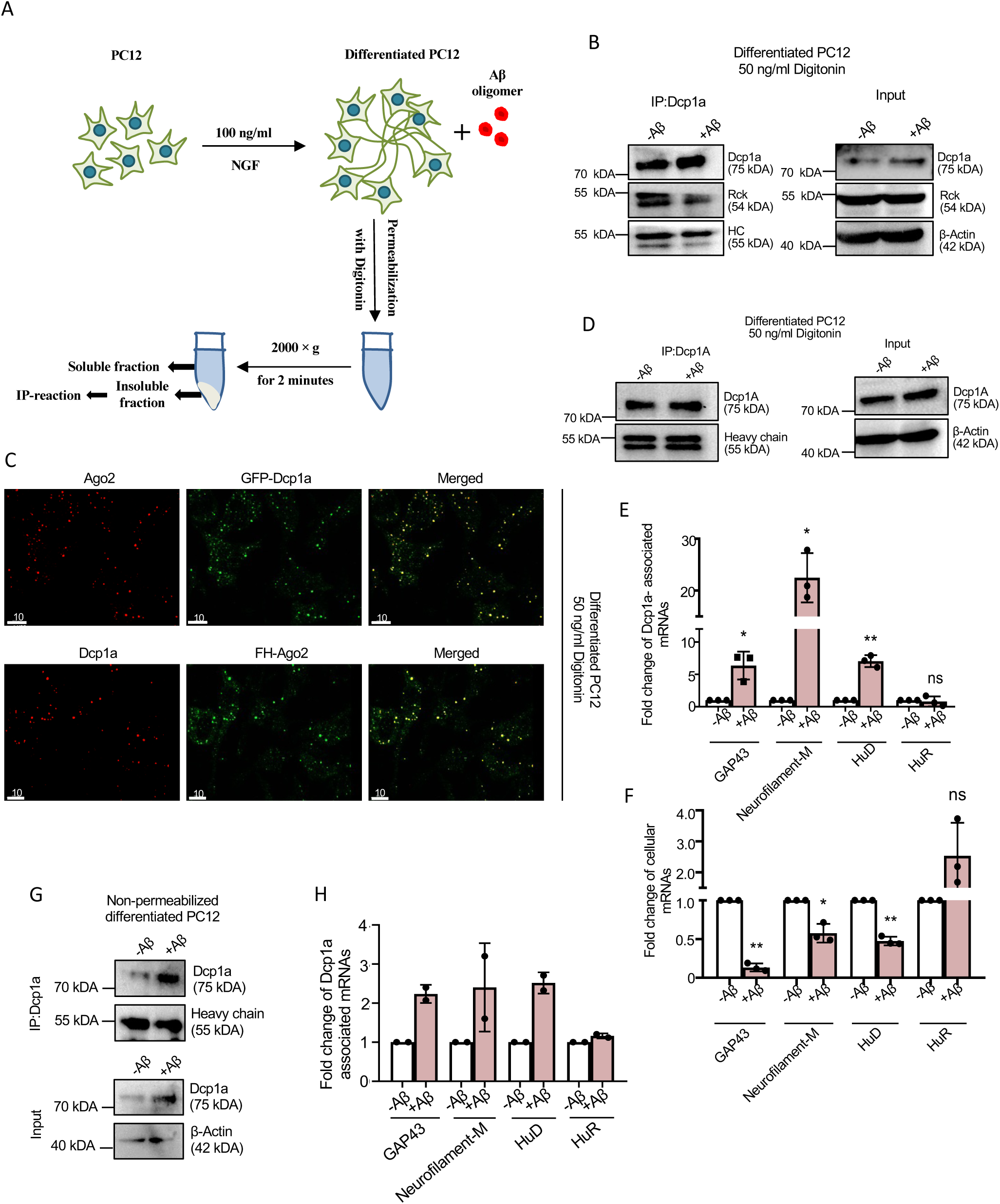
Altered association of mRNAs with PB-associated Dcp1a in amyloid beta oligomer-treated differentiated PC12 cells. **(A)** Schematic representation showing separation of soluble and non-soluble fractions from Aβ _(1-42)_ treated differentiated PC12 cells permeabilized with Digitonin. **(B)** Immunoprecipitation of endogenous Dcp1a protein from 0 or 2.5 µM Aβ _(1-42)_ treated (24 hours) differentiated PC12 cells permeabilized with 50 ng/mL Digitonin. Western blot images of pulled-down Dcp1a and associated RCK/p54 levels in Dcp1a immunoprecipitated materials isolated from extract made with Digitonin-permeabilized and PB-enriched ghost NGF-differentiated PC12 cells. Dcp1a and RCK/p54 levels, along with β-Actin as a loading control, are shown in the inputs used for immunoprecipitation. Molecular weight markers are indicated. HC: heavy chain. **(C)** Confocal Images showing co-localization between exogenously expressed GFP-Dcp1a (Green) and endogenous Ago2 (Red) in 72-hour differentiated (with 100 ng/mL NGF) PC12 cells permeabilized with 50 ng/mL Digitonin (Upper panel). Confocal Images showing co-localization between exogenously expressed FH-Ago2 (Green) and endogenous Dcp1a (Red) in 72-hour differentiated PC12 cells permeabilized with Digitonin (lower panel). Merged images are shown. Fields have been detected in 63X magnification. Scale bar represents 10 µm. **(D-E)** Immunoprecipitation of endogenous Dcp1a protein from 0 or 2.5 µM Aβ _(1-42)_ treated (24 hours) differentiated PC12 cells permeabilized with 50 ng/mL Digitonin. Western blot images of the pulled-down Dcp1a levels and input Dcp1a levels, with β-Actin as a loading control, are shown. Molecular weight markers are indicated. HC: Heavy chain (D). qRT-PCR based quantification of pulled-down Dcp1a associated GAP43, Neurofilament-M, HuD and HuR mRNA levels normalized against immunoprecipitated Dcp1a levels [p=0.0071 (for GAP43), p=0.0120 (for Neurofilament-M), p=0.0495 (for HuD), p=0.0490 (for HuR), n=3 independent experiments] (E). **(F)** qRT-PCR based quantification of cellular GAP43, HuD and HuR mRNA levels normalized against GAPDH level in 0 or 2.5 µM Aβ _(1-42)_ treated (24 hours) differentiated PC12 cells permeabilized with 50 ng/mL Digitonin [p=0.0254 (for GAP43), p=0.0352 (for HuD), p=0.1195 (for HuR), n=3 independent experiments]. Molecular weight markers are shown. Data represents means ± SDs; ns, non-significant, *p < 0.05, **p< 0.01, ***p < 0.001, ****p < 0.0001. p values were obtained by using two-tailed paired Student’s t test. **(G-H)** Immunoprecipitation of total endogenous Dcp1a protein from both 0 or 2.5 µM Aβ _(1-42)_ treated differentiated PC12 cells without digitonin permeabilization. Western blot images of the pulled-down Dcp1a level and input Dcp1a levels, with β-Actin as a loading control, are shown. Molecular weight markers are indicated. HC: Heavy Chain (G). qRT-PCR-based quantification of associated GAP43, Neurofilament-M, HuD, and HuR mRNA levels normalized against relative levels of Dcp1a immunoprecipitated (n=2 independent experiments) (H).

Consequently, total cellular levels of these mRNAs decreased upon Aβ(1-42) exposure, contrary to their typical upregulation during differentiation (**Fig. 2F**). This reduction implies that Aβ(1-42) may impair the efficient translation of these key mRNAs, thereby disrupting the differentiation process. Notably, a similar pattern of increased Dcp1a association with GAP-43, Neurofilament-M, and HuD mRNAs was observed when Dcp1a immunoprecipitation was performed from total cell lysates without prior digitonin permeabilization to reduce cytoplasmic noise. However, the total cellular Dcp1a enrichment of GAP-43, HuD, and Neurofilament-M mRNAs was much less compared to mRNAs associated with Dcp1a isolated from PB-enriched materials (**Fig. 2 G and H**). Thus, PB enrichment found to be helpful in reducing noise and enhancing signals of PB-localized mRNAs for PB-specific proteins like Dcp1a immunoprecipitated, enabling the identification of mRNAs altered in PBs caused by amyloid beta in NGF-differentiated PC12 cells.

### Amyloid beta oligomers affect mRNA association with PBs in glioblastoma cells

In glial cells treated with Aβ_(1-42)_, elevated mRNA levels of pro-inflammatory cytokines have been reported, linked to disrupted repressive microRNA (miRNA) function caused by retention of miRNPs in endosomes with impaired lysosomal targeting and recycling (De and Bhattacharyya, 2021; De et al., 2021). To investigate the association of these cytokine mRNAs with processing bodies (PBs) upon exposure of glioblastoma cells to amyloidogenic oligomers, we used the detergent-based permeabilization protocol to isolate phase-separated, non-soluble PBs-associated Dcp1a from C6 astroglial cells (**Fig. 3A**). After optimizing digitonin concentration for this assay in C6 cells, we confirmed that 50 ng/mL digitonin efficiently isolated PB-associated Dcp1a bodies, consistent with prior optimization results with PC12 cells (**Fig. 3B-C, Fig. S2A**).

**Figure 3.**
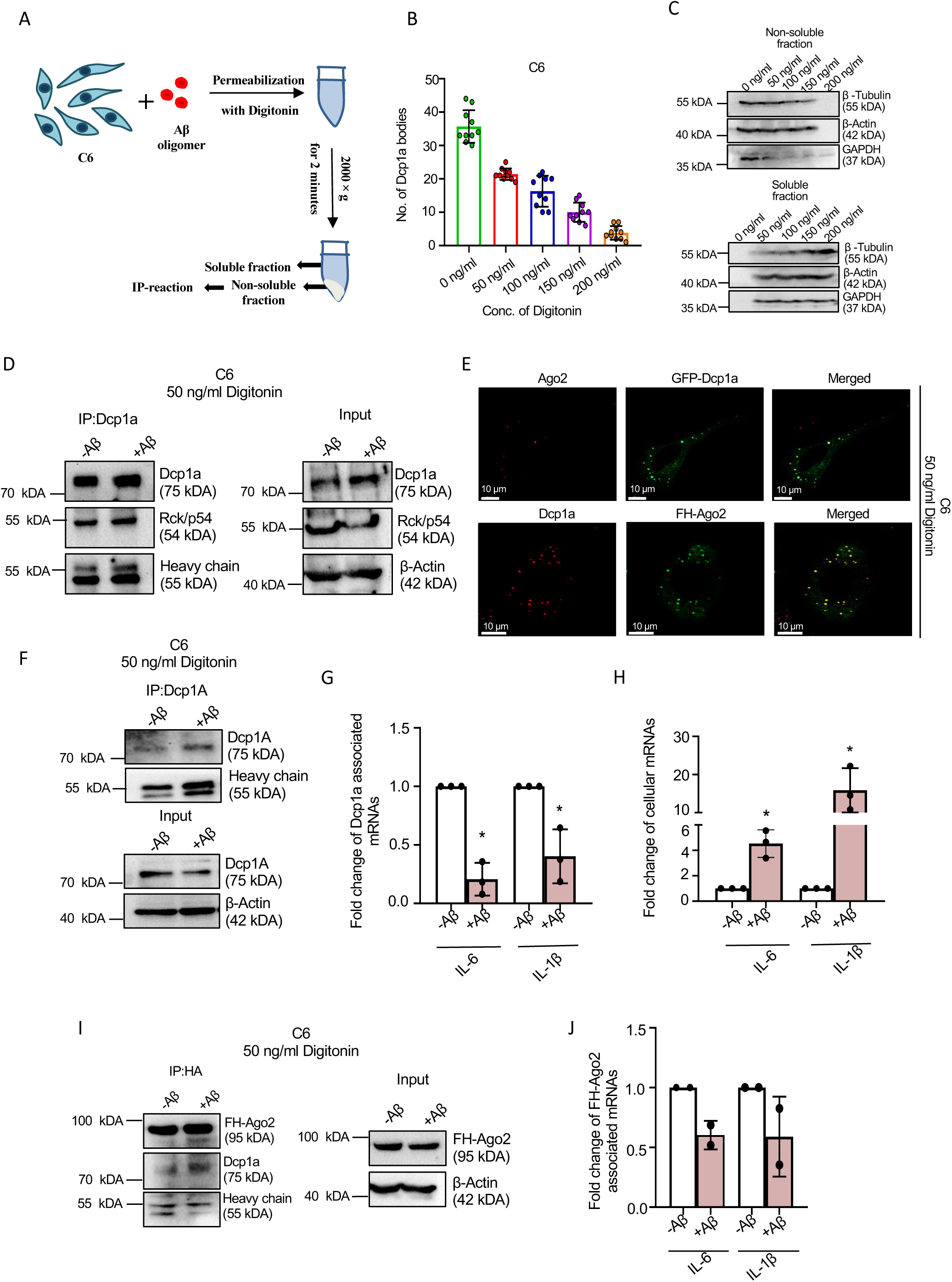
Changed pattern of mRNA association with PB-localized Dcp1a in amyloid beta-treated astroglial cells. **(A)** Schematic representation showing separation of soluble and non-soluble fractions from amyloid beta oligomer-treated C6 cells permeabilized with Digitonin. **(B)** Quantification of GFP-Dcp1a bodies in GFP-Dcp1a expressing C6 cells permeabilized with increasing concentration of Digitonin (0, 50, 100, 150, 200 ng/mL) for 10 minutes at 4°C. **(C)** Western blot of endogenous β-Tubulin, β-Actin and GAPDH were done to measure the expression levels of these proteins in the non-soluble fraction (upper panel) and soluble fraction (lower panel) of C6 cells permeabilized with increasing concentration of Digitonin (0, 50, 100, 150, 200 ng/ml). Molecular weight markers are shown. **(D)** Immunoprecipitation of endogenous Dcp1a protein from both 0 or 2.5 µM Aβ _(1-42)_ treated (24 hours) C6 cells permeabilized with 50 ng/ml Digitonin. Western blot images of immunoprecipitated Dcp1a and associated RCK/p54 levels, and input Dcp1a and RCK/p54 levels with β-Actin as loading control are shown. Molecular weight markers are also shown. HC: Heavy Chain. **(E)** Confocal Images showing co-localization between exogenously expressed GFP-Dcp1a (Green) and endogenous Ago2 (Red) in C6 cells permeabilized with 50 ng/mL Digitonin (Upper panel). Confocal Images showing co-localization between exogenously expressed FH-Ago2 (Green) and endogenous Dcp1a (Red) in C6 cells permeabilized with 50 ng/mL Digitonin (lower panel). Merged images are shown. Fields have been detected in 63X magnification. Scale bar represents 10 µm. **(F-G)** Immunoprecipitation of endogenous Dcp1a protein from both 0 or 2.5 µM Aβ _(1-42)_ treated C6 cells permeabilized with 50 ng/ml Digitonin. Western blot images of immunoprecipitated Dcp1a level, and input Dcp1a level with β-Actin as loading control, are shown. Molecular weight markers are also shown, HC: Heavy Chain (F). qRT-PCR-based quantification of immunoprecipitated Dcp1a-associated IL-6 and IL-1β mRNA levels normalized against immunoprecipitated Dcp1a levels [(p=0.0102 (for IL-6), p=0.0463 (for IL-1β), n=3 independent experiments)] (G). Data represents means ± SDs; ns, non-significant, *p < 0.05, **p< 0.01, ***p < 0.001, ****p < 0.0001. p-values were obtained by using a two-tailed paired Student’s t-test. **(H)** qRT-PCR-based quantification of cellular IL-6 and IL-1β mRNA levels normalized against GAPDH level in 0 or 2.5 µM Aβ _(1-42)_ treated C6 cells permeabilized with 50 ng/mL Digitonin. [(p=0.0301 (for IL-6), p=0.0472 (for IL-1β), n=3 independent experiments)]. Data represents means ± SDs; ns, non-significant, *p < 0.05, **p< 0.01, ***p < 0.001, ****p < 0.0001. p-values were obtained by using a two-tailed paired Student’s t-test. **(I-J)** Immunoprecipitation of exogenously expressed FH-Ago2 protein by anti-HA antibody from both 0 or 2.5 µM Aβ _(1-42)_ treated C6 cells permeabilized with 50 ng/ml Digitonin. Western blot images of immunoprecipitated FH-Ago2 and associated Dcp1a levels, and input FH-Ago2 level with β-Actin as loading control are shown. Molecular weight markers are shown. HC: Heavy Chain (I). qRT-PCR based quantification of HA-Ago2 associated IL-6 and IL-1β mRNA levels normalized against relative pulled down FH-Ago2 levels (n=2 independent experiments) (J).

We immunoprecipitated Dcp1a from the non-soluble fraction of digitonin-permeabilized C6 cells, both treated with and without Aβ(1-42), followed by RT-PCR quantification of PB-associated pro-inflammatory cytokine mRNAs, specifically IL-6 and IL-1β. The immunoprecipitated Dcp1a was found to be associated with RCK/p64, a PB marker, and confocal imaging revealed that Ago2 was also located with Dcp1a after digitonin-permeabilization of C6 cells (**Fig. 3D-E**). As expected, total cellular levels of IL-6 and IL-1β mRNAs increased after Aβ_(1-42)_ treatment, consistent with previous findings. However, their association with PBs was significantly reduced, indicating a release from translational repression and an increase in translational activity in the presence of amyloid oligomers (**Fig. 3F-H**). A similar decrease in PB association was observed when total Dcp1a was immunoprecipitated from non-permeabilized C6 cells (**Fig. S2B-C**).

Ago2 is found to colocalize with PBs in control and A-treated cells, and we analysed the interaction between FH-Ago2 and cytokine mRNAs in detergent-permeabilized FH-Ago2-expressing C6 cells. Immunoprecipitation assays revealed a significant reduction in the association of IL-6 and IL-1β mRNA with Ago2 following Aβ treatment, indicating that amyloid oligomers disrupt Ago2-mRNA interactions, which may contribute to the increased cytokine expression observed (**Fig. 3H and J**). Additionally, FH-Ago2 proteins co-immunoprecipitated with Dcp1a (**Fig. 3I**) and co-localized with Dcp1a in immunofluorescence assays (**Fig. 3E**), supporting the interplay between these PB components in cytokine mRNA regulation and confirming the effectiveness of this technique with multiple PB components (i.e., Dcp1a and Ago2).

### Validation of biochemical data using *In Situ Hybridization*

In parallel, to confirm the changes in the PB association of specific pro-inflammatory cytokine mRNAs, we used RNA fluorescence in situ hybridization (RNA-FISH) techniques to verify our biochemical data(Pillai et al., 2004). C6 cells were co-transfected with plasmids encoding GFP-tagged Dcp1a and reporter constructs expressing Renilla luciferase (RL) fused to either IL-6 or IL-1β mRNA sequences (RL-IL-6 and RL-IL-1β). After permeabilization with 50 ng/mL digitonin, we applied Cy3-labeled oligonucleotide probes complementary to the RL sequence to specifically detect these reporter mRNAs in C6 cells (**Fig. 4A**).

**Figure 4.**
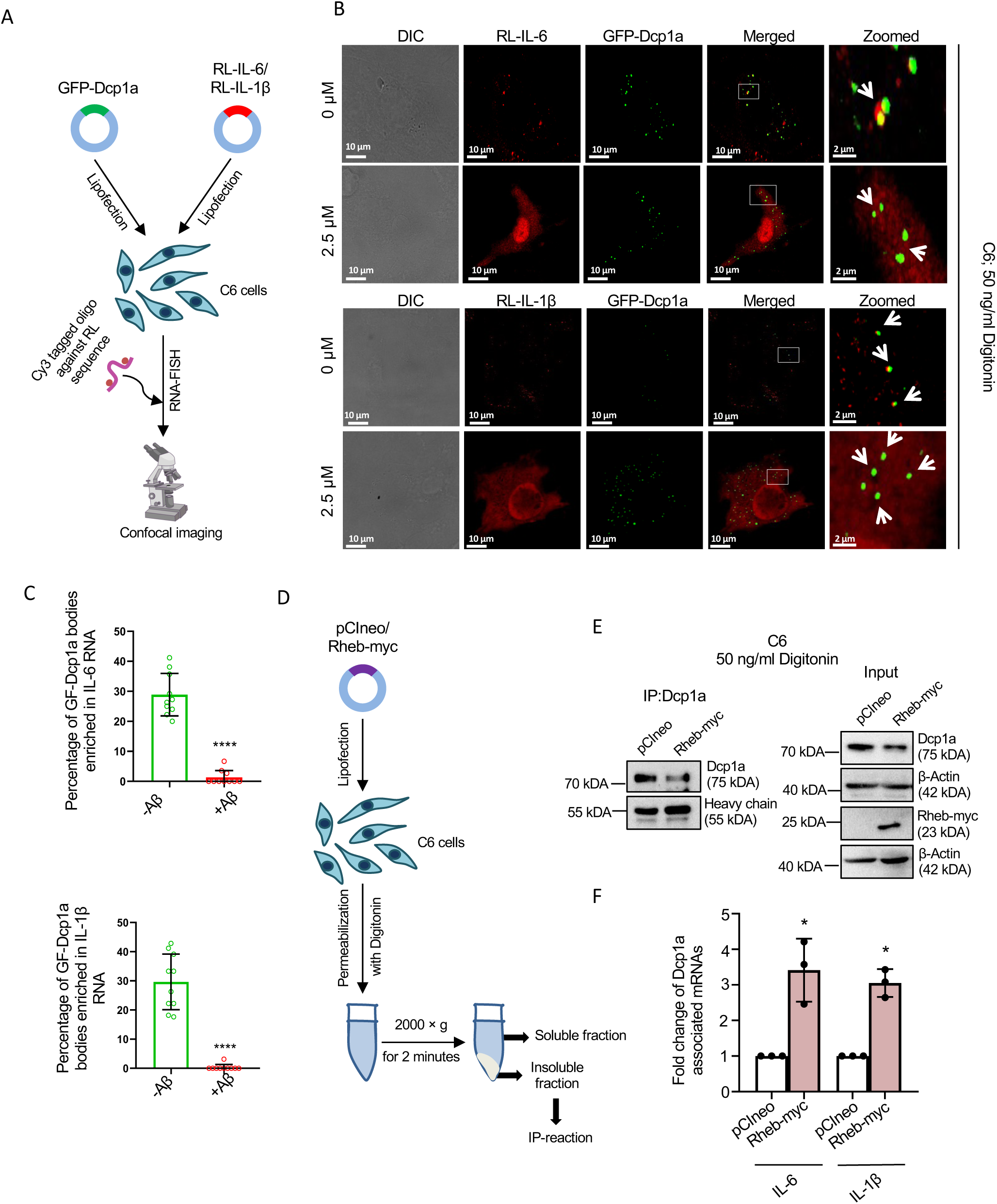
Rescue in Dcp1a-association and PB-localization of cytokine mRNAs by Rheb-Myc in amyloid-beta treated C6 cells. **(A)** Schematic representation of RNA-FISH experiment using Cy3-tagged oligos against RL sequence in GFP-Dcp1a, and RL-IL-6, or RL-IL-1β co-transfected amyloid beta oligomer-treated C6 cells. **(B)** Confocal Images showing co-localization between exogenously expressed GFP-Dcp1a (Green) and RL-IL-6 mRNAs (Red) by RNA-FISH in 0 or 2.5 µM Aβ _(1-42)_ treated C6 cells (Upper panel). Confocal Images showing co-localization between exogenously expressed GFP-Dcp1a (Green) and RL-IL-1β mRNAs (Red) by RNA-FISH in 0 or 2.5 µM Aβ _(1-42)_ treated C6 cells (lower panel). Merged images are shown. Fields have been detected in 63X magnification. Scale bar of non-zoomed images represents 10 µm, whereas those of zoomed images represent 2 µm. **(C)** Quantification of the percentage of GFP-Dcp1a, as in experiments described in panel B, positive for RL-reporters (n=10). **(D)** Schematic representation showing separation of soluble and non-soluble fractions from Aβ _(1-42)_ treated pCIneo or Rheb-myc expressing C6 cells permeabilized with Digitonin. **(E-F)** Immunoprecipitation of endogenous Dcp1a protein from both pCIneo or Rheb-Myc expressing 2.5 µM Aβ _(1-42)_ treated C6 cells permeabilized with 50 ng/ml Digitonin. Western blot images of immunoprecipitated Dcp1a levels and input Dcp1a and Rheb-Myc levels, with β-Actin as a loading control, are shown. Molecular weight markers are also shown. HC: Heavy Chain (E). qRT-PCR based quantification of associated IL-6 and IL-1β mRNA levels normalized against relative pulled down Dcp1a levels [(p=0.0421 (for IL-6), p=0.0119 (for IL-1β), n=3 independent experiments] (F). Data represents means ± SDs; ns, non-significant, *p < 0.05, **p< 0.01, ***p < 0.001, ****p < 0.0001. p-values were obtained by using a two-tailed paired Student’s t-test

Image analysis showed a significant increase in RL-IL-6 and RL-IL-1β mRNA signal intensity in cells treated with Aβ oligomers compared to untreated controls, indicating increased expression or stabilization of these cytokine transcripts under amyloidogenic conditions. Notably, despite this rise in mRNA levels, there was a decrease in the co-localization of RL-tagged cytokine mRNAs with GFP-Dcp1a-positive PBs. This suggests that Aβ oligomers interfere with the sequestration of excess cytokine mRNAs within PBs, possibly allowing their release from translational repression (**Fig. 4B-C**). These RNA-FISH results support our immunoprecipitation data, collectively indicating that amyloidogenic oligomers disrupt the regular regulation of cytokine mRNAs by PBs, contributing to their elevated expression. This dysregulation may partly drive the inflammatory response observed in glial cells exposed to stress induced by amyloid proteins, establishing a mechanistic link between amyloid pathology and the post-transcriptional regulation of inflammatory mediators. Interestingly, Rheb-Myc, the activator of mTORC1, has been reported to rescue cytokine repression in Aβ-treated cells(De et al., 2021). In the Dcp1a-IP reaction performed with Rheb-Myc-expressing cells treated with Aβ oligomers, we observed a rescue of Dcp1a association with IL-1β and IL-6, confirming the reciprocal relationship between the PB-localization of cytokine mRNAs and the expression of Rheb-Myc proteins. This rescue, mediated by Rheb-Myc, an activator of mTORC1, underscores how amyloid treatment affects cytokine mRNA regulation (**Fig. 4D-F**).

## Discussion

RNA processing bodies (P-bodies or PBs) are specialized, dynamic cytoplasmic granules that play crucial roles in the post-transcriptional regulation of gene expression. These structures are formed through liquid-liquid phase separation, a process in which proteins and RNA molecules interact to create dense, membrane-less compartments. Phase separation is driven by complex networks of protein interactions involved in mRNA decay, translational repression, and RNA surveillance. Although PBs are primarily known for their role in mRNA degradation, they also serve as storage sites for mRNAs that are translationally repressed (Decker and Parker, 2012; Jain and Parker, 2013; Parker and Sheth, 2007).

By sequestering mRNAs in a translationally inactive state, PBs regulate the spatial and temporal dynamics of protein production within the cytoplasm. This regulation ensures that proteins are synthesized at the correct time and in the appropriate quantities, particularly when cells encounter fluctuating environmental or developmental signals. For example, cells must adjust protein production in response to stress, changes in nutrition, or differentiation cues, with PBs acting as key regulators of these adaptive processes (Bhattacharyya et al., 2006; Cougot et al., 2008).

In neurons, the regulation of mRNA translation is crucial, as these cells must maintain a balance between undifferentiated and differentiated states. Neurons exhibit intricate morphological features and functional states that rely on the precise control of mRNA translation, both locally at synapses and in perinuclear regions within the cell. The local translation of mRNAs is crucial for synaptic plasticity, learning, memory, and neuronal development (Holt et al., 2019). Any disruption in PB function or loss of mRNA translation control can lead to severe neurological disorders, such as Alzheimer’s disease, Parkinson’s disease, or Huntington’s disease (Li and Sun, 2025). Thus, understanding how mRNAs are processed and localized in PBs under different cellular conditions may offer insights into the molecular mechanisms underlying these diseases.

A significant advancement in PB research was made by Hubstenberger et al. in 2017, who utilized Fluorescence-Activated Particle Sorting (FAPS) to purify PBs from human epithelial cells. This high-throughput technique allowed for the identification of hundreds of proteins and RNA molecules that comprise PBs, providing a detailed view of these RNA-protein granules (Hubstenberger et al., 2017). However, FAPS has its limitations, including the need for specialized equipment such as high-throughput fluorescence sorters and the potential for introducing artifacts during sorting and purification. These constraints can hinder the complete characterization of PBs under various experimental conditions, particularly in settings with limited access to advanced instruments.

To overcome these challenges, we developed a more accessible and flexible method for isolating PBs. Our approach employs a detergent-based technique to separate the soluble and non-soluble fractions of the cytoplasm, followed by solubilization, and removal of insoluble membranes by centrifugation to enrich and isolate PBs. We selected digitonin as the detergent due to its ability to selectively permeabilize the plasma membrane in a controlled manner, thereby maintaining cellular integrity and preserving PB structure (Adam, 2016). The non-soluble, phase-separated fraction, which contains PBs, is then enriched by centrifugation. To further isolate PBs, we conducted immunoprecipitation using an antibody against Dcp1a, a marker protein of PBs that is highly abundant in these structures. This step enables the selective capture of PBs and their associated mRNAs.

Once isolated, we analysed the mRNA content of PBs using quantitative reverse-transcription polymerase chain reaction (qRT-PCR). This method enabled us to examine the specific mRNAs associated with PBs quantitatively under various experimental conditions, providing insights into the regulatory processes within these granules. We initially focused on mRNAs involved in neuronal differentiation, given their importance in neuronal function and development. By tracking the PB-association status of these mRNAs, we assessed how their localization changes during differentiation and explored the role of PBs in this process.

Next, we investigated how amyloidogenic proteins, such as amyloid-beta, influence mRNA localization within PBs during neuronal differentiation. The aggregation of amyloidogenic proteins is a hallmark of neurodegenerative diseases, and their accumulation disrupts normal cellular processes, including mRNA regulation and translation(De and Bhattacharyya, 2021; De et al., 2021). Understanding how amyloidogenic proteins impact PB function and mRNA localization could offer valuable insights into the molecular mechanisms of neurodegeneration(Ray et al., 2025). We also extended our study to glial cells, which play a crucial role in inflammation during neurodegenerative diseases. We examined how amyloid-induced inflammation reduces the localization of pro-inflammatory cytokine mRNAs in PBs.

To confirm the identity of the Dcp1a-positive granules as PBs, we performed co-localization studies with other established PB marker proteins, such as Ago2 and Rck/p54. The co-localization of these additional marker proteins within the same granules further validated our characterization of the isolated structures as authentic PBs. By using this method across neuronal and glial cell lines, we gained a comprehensive understanding of PB dynamics in different cell types and under varying conditions.

Post-transcriptional modifications, such as mRNA degradation, translation, and sequestration in PBs, are essential for maintaining cellular homeostasis. Disruptions in these processes can lead to a wide range of diseases, including neurological disorders, cancer, and inflammatory conditions. Our study demonstrates that the method we developed for isolating and characterizing PBs can be widely applied to investigate mRNA association with PBs in various cellular and physiological contexts. This approach holds significant potential for uncovering how PBs contribute to disease pathogenesis, opening new avenues for therapeutic strategies. Moving forward, we anticipate that this accessible and adaptable method will become a valuable tool for studying the molecular mechanisms and importance of Phase separation as a key post-transcriptional regulation happening in health and disease.

### Limitations of the study

While our detergent-based method for isolating PBs offers a practical and accessible alternative to high-throughput approaches like FAPS, it is not without limitations. PBs are inherently heterogeneous in their composition, and distinct subpopulations likely exist within a single cell, each potentially involved in different aspects of mRNA regulation(Luo et al., 2018). Our assay, which relies on immunoprecipitation using the PB marker Dcp1a, captures only a representative subset of these structures. As such, the isolated PBs may not fully reflect the diversity of PB populations or their complete RNA content. Additionally, while our co-localization with known PB markers supports the authenticity of these granules as “traditional” PBs, functional heterogeneity and context-specific roles of PBs may still be underrepresented. Future refinements, including multi-marker approaches and single-particle analyses, may help uncover the full complexity of PB biology.

### Experimental Procedures

## EXPERIMENTAL MODEL AND STUDY PARTICIPANT DETAILS

PC12 cells were cultured in Dulbecco’s modified Eagle’s medium (DMEM; GIBCO) supplemented with 10% heat-inactivated horse serum (HS; GIBCO), 5% heat-inactivated fetal bovine serum (FBS; GIBCO) and 1% Penicillin-Streptomycin. PC12 cells were differentiated by using 100 ng/ml NGF (Promega) added in differentiation medium containing Dulbecco’s modified Eagle’s medium (DMEM; GIBCO) supplemented with 0.75% heat-inactivated horse serum (HS; GIBCO), 0.25% heat-inactivated fetal bovine serum (FBS; GIBCO) for 72 hours. C6 cells were cultured in Dulbecco’s modified Eagle’s medium (DMEM; GIBCO) supplemented with 10% heat-inactivated fetal bovine serum (FBS; GIBCO) and 1% Penicillin-Streptomycin. Cells were treated with Amyloid oligomers at a strength of 2.5 µM from a stock of 200 µM for 24 hours.

## METHOD DETAILS

### Cell transfection

All the transfections were done using Lipofectamine 2000 reagent (Invitrogen) following the manufacturer’s protocol. For immunoprecipitation, cells were transfected with 2 µg of plasmids in a 60 mm format. For immunofluorescence and RNA-FISH, cells were transfected with 500 ng of plasmids in a 12-well format. Transfections were done for 6 hours for both cell lines followed by a media change. Plasmid transfected cells were then split to a larger growing surface after 24 hours of transfection.

### P-Body isolation

Confluent cells (∼6 × 10L) cultured in a 90 mm dish were washed with cold 1× PBS (pH 7.4), harvested by scraping, and pelleted by centrifugation at 800 ×Lg for 5 minutes at 4L°C. The cell pellet was resuspended in 500 µL of freshly prepared permeabilization buffer containing 50Lng/mL Digitonin in PBS and incubated for 10 minutes at 4L°C with gentle rotation. Subsequent centrifugation at 2000L×Lg for 2 minutes at 4L°C yielded the soluble fraction in the supernatant and the non-soluble components in the pellet. The pellet was washed with PBS and re-centrifuged to remove residual Digitonin. For P-body isolation, 30 µL of Protein G agarose beads per sample were washed with IP buffer (20 mM Tris-HCl pH 7.5, 150 mM KCl, 5 mM MgCl₂, 1 mM DTT, 1× PMSF, and RNase inhibitor), blocked with 5% BSA in lysis buffer for 1 hour at 4L°C, and incubated with Dcp1a antibody (1:100 dilution) in lysis buffer for 4 hours at 4 °C with rotation. The non-soluble pellet was lysed in 350 µL of lysis buffer. For C6 cells, the lysate was sonicated (3 cycles of 10 pulses each at 1 pulse per second) and centrifuged at 16,000 ×Lg for 10 minutes at 4 °C. For PC12 cells, sonication was omitted, and the lysate was directly centrifuged at 3000 ×Lg for 10 minutes at 4 °C. The cleared lysate was incubated overnight at 4 °C with the antibody-bound beads. The following day, the beads were washed and divided for downstream protein (western blot) and RNA (qRT-PCR) analyses.

### RNA isolation and qRT-PCR

Total RNA was isolated by using TRIzol or TRIzol LS reagent (Invitrogen) according to the manufacturer’s protocol. mRNA levels were measured by two step Real-time qRT-PCR method using Eurogentec Reverse Transcriptase Core Kit and MESA GREEN qPCR Master Mix Plus for SYBR Assay. 100 to 200 ng of RNA was used for detection of cellular mRNA levels and equal volume of RNA was used to analyze Dcp1a associated mRNA levels. GAPDH was used as endogenous control. All the PCRs were done in 7500 Applied Biosystem Real-Time system. The Reverse Transcription reaction conditions were 25 °C for 10 minutes, 48 °C for 30 minutes, 95 °C for 5 minutes and finally product held at 4 °C. The qPCR conditions were 95 °C for 5 minutes followed by 40 cycles of 95 °C for 15 seconds, 60°C for 1 minute.

### Western Blot

The samples were diluted in 5X sample loading buffer containing 312.5 mM Tris-HCl pH 6.8, 10% SDS, 50% glycerol, 250 mM DTT, 0.5% Bromophenol blue and heated at 95°C for 10 minutes. Following SDS-polyacrylamide gel electrophoresis, proteins were transferred to PVDF membranes (Millipore). Membranes were then blocked using 1X TBS (Tris Buffered Saline) supplemented with 0.1% Tween-20 (1X TBST) and 3% BSA (HiMedia). Primary antibodies were added in 1X TBST containing 3% BSA for around 16 hours at 4°C. After overnight incubation with antibody, the membranes were washed thrice for 5 minutes each with 1X TBST at room temperature to remove excess primary antibodies. It is then followed by incubation of membranes at room temperature for 1-1.5 hours with secondary antibodies conjugated with horseradish peroxidase (1:8000 dilution) in 3% BSA containing 1X TBST. Excess secondary antibodies were washed thrice with 1X TBST at room temperature. Antigen-antibody complexes were detected with West Pico Chemiluminescent, Luminata Forte Western HRP substrate or West Femto Maximum Sensitivity substrates using standard manufacturer’s protocol. Imaging of all western blots was done using a UVP BioImager 600 system equipped with Vision Works Life Science software (UVP) V6.80.

### Immunofluorescence

Cells grown on coverslips were fixed with 4% Paraformaldehyde dissolved in 1X Phosphate Buffered Saline (PBS) at room temperature for 30 minutes in the dark. Coverslips were then washed thrice with 1X PBS for 5 minutes each, which was followed by blocking and permeabilization with 1X PBS containing 10% goat serum (Gibco), 20% bovine serum albumin (HiMedia), and 0.1% Triton X-100 (CALBIOCHEM) for 30 minutes at room temperature. After being washed thrice with 1X PBS for 5 minutes each, Coverslips were then incubated with Primary antibody, diluted in 1× PBS with 20% BSA and kept overnight in a humid chamber at 4°C. Following three washes with 1X PBS for 5 minutes each, secondary antibody labelled with Alexa Fluor diluted in 1X PBS containing 20% BSA, was added on coverslips and the incubation was done in a humid chamber for 1 hour at room temperature in the dark. To remove excess secondary antibodies, the coverslips were washed thrice with 1X PBS for 5 minutes each and mounted with Vectashield mounting medium with DAPI (4′,6-diamidino-2-phenylindole; Vector).

### Cy3 labelling of oligos

30 µl of DMSO containing 1 vial of CY3 monoreactive dye (GE HealthCare) was mixed with 10 µg of oligos against RL sequences re-suspended in 0.1 M NaHCO_3_ buffer (pH 8.8) and kept for 24 hours at room temperature in the dark. To remove unreacted dye, it was followed by two rounds of Ethanol precipitations and 3 rounds of purifications using miniQuick Spin Oligo Columns (Roche). The oligos were obtained from Eurogentec having sequences as follows-

RL 1: 5’aT*cacaaagatgatT*ttctttggaaggtT*ca;

RL 2: 5’aT*tagctggaggcagcgT*taccatgcagaaa;

RL 3: 5’aT*agtccagcacgtT*catttgcttgcagT*ga.

### RNA-FISH (Fluorescence in-situ hybridization)

Cells grown on coverslips were fixed with 4% paraformaldehyde dissolved in 1X Phosphate Buffered Saline (PBS) at room temperature for 30 minutes in the dark. Following three washes with 1X PBS for 5 minutes each, the coverslips were permeabilized with 70% alcohol overnight at 4 °C. Cells were then rehydrated with 25% Formamide containing 2X SSC at room temperature for 10 minutes and incubated with a hybridization solution containing 10% Dextran Sulphate, 2mM Vanadyl Ribonucleoside Complex, 0.02% RNase-free BSA, 40 µg Salmon Sperm DNA, 2X SSC, 25% Formamide, 30 ng probe at 37 °C overnight in a humid chamber. After being washed twice with 2X SSC containing 20% Formamide for 15 minutes each at 37 °C, coverslips were mounted with Vectashield mounting medium with DAPI (4′,6-diamidino-2-phenylindole; Vector).

### Preparation of Amyloid beta

100% 1,1,1,3,3,3, hexafluoro-2-propanol (HFIP) was used to solubilize lyophilized Aβ _(1-42)_ and Aβ _(1-40)_ (American Peptide) to a final stock concentration of 1 mM. HFIP was then removed by evaporation using SpeedVac (Eppendorf). The pellet was solubilized in anhydrous DMSO followed by sonication in a bath sonicator for 40 minutes at 37 °C. The final stock solution was made at a concentration of 5 mM and stored at -80 °C. For preparing oligomers of both Aβ _(1-42)_ and Aβ _(1-40)_ the stock peptides were diluted in 1X Phosphate Buffered Saline (PBS) and Sodium Dodecyl Sulphate (SDS) at a concentration of 0.2% to make the concentration of the peptide 400 µM. The solution was then incubated overnight at 37 °C in a rotating motion. it was then further diluted with 1X PBS to a final concentration of 200 µM and again incubated overnight at 37 °C for proper oligomerization before use.

## QUANTIFICATION AND STATISTICAL ANALYSIS

### Image capture and post-capture image processing

All the images were captured using Zeiss LSM800 confocal microscope and Leica confocal microscope SP8 (Leica Microscope Systems; Wetzler, Germany). All image processing were done using Imaris7 software developed by BITPLANE AG Scientific software.

### Statistical analysis

All graphs and statistical analyses were generated using Prism (v5.00 and v8.00) software (GraphPad, San Diego, CA). Nonparametric two-tailed paired *t-*tests were used for analysis. *P* values of <0.05 were considered statistically significant, and *P* values of >0.05 were not significant. Error bars indicate means ± standard deviation (SDs).

### Supplemental information

Information on plasmids, oligos, antibodies, and primers is available in Supplementary Tables S1 to S4.

### Resource availability

All resources reported in this manuscript are available in the data presented within the manuscript figures and resource tables.

### Lead contact

Further information and requests for resources and reagents should be directed to and will be fulfilled by the lead contact, Dr. Suvendra N. Bhattacharyya.

### Materials availability

This study did not produce any unique reagents.

### Data and Code Availability

- All data supporting the findings of this study are available from the corresponding authors upon request.
- This paper does not report the original code.
- Any additional information required to reanalyse the data reported in this paper is available from the lead contact upon request.

## Acknowledgments

We thank Witold Filipowicz, Edouard Bertrand, and Gunter Meister for the plasmid constructs used in this study. We also thank the Council for Scientific and Industrial Research (CSIR) for the SR. SNB was supported by the Swarnajayanti Fellowship (DST/SJF/LSA-03/2014-15) from the Department of Science and Technology, Govt. of India. The work received support from a High-Risk High-Reward Grant (HRR/2016/000093) from the same department, as well as a Fund from CEFIPRA. SNB is supported by the Start-Up Support Grant from the University of Nebraska, USA, and K.M. acknowledges the Lieberman Research Award from the Department of Anesthesiology, UNMC.

## Author contributions

Sritama Ray: Data Curation, Formal Analysis, Investigation, Methodology. Kamalika Mukherjee: Data Curation, Formal Analysis, Investigation, Writing-Review and Editing, Conceptualization, Methodology, Supervision, Writing-Review and Editing. Suvendra N Bhattacharyya: Data Curation, Formal Analysis, Investigation, Writing-Review and Editing, Conceptualization, Funding Acquisition, Methodology, Supervision.

