## Supplementary Info for "Isolation and Quantification of mRNAs from Subcellular Phase-Separated Structures from Detergent Permeabilized Brain Cells"

**Supplementary figure and legends**

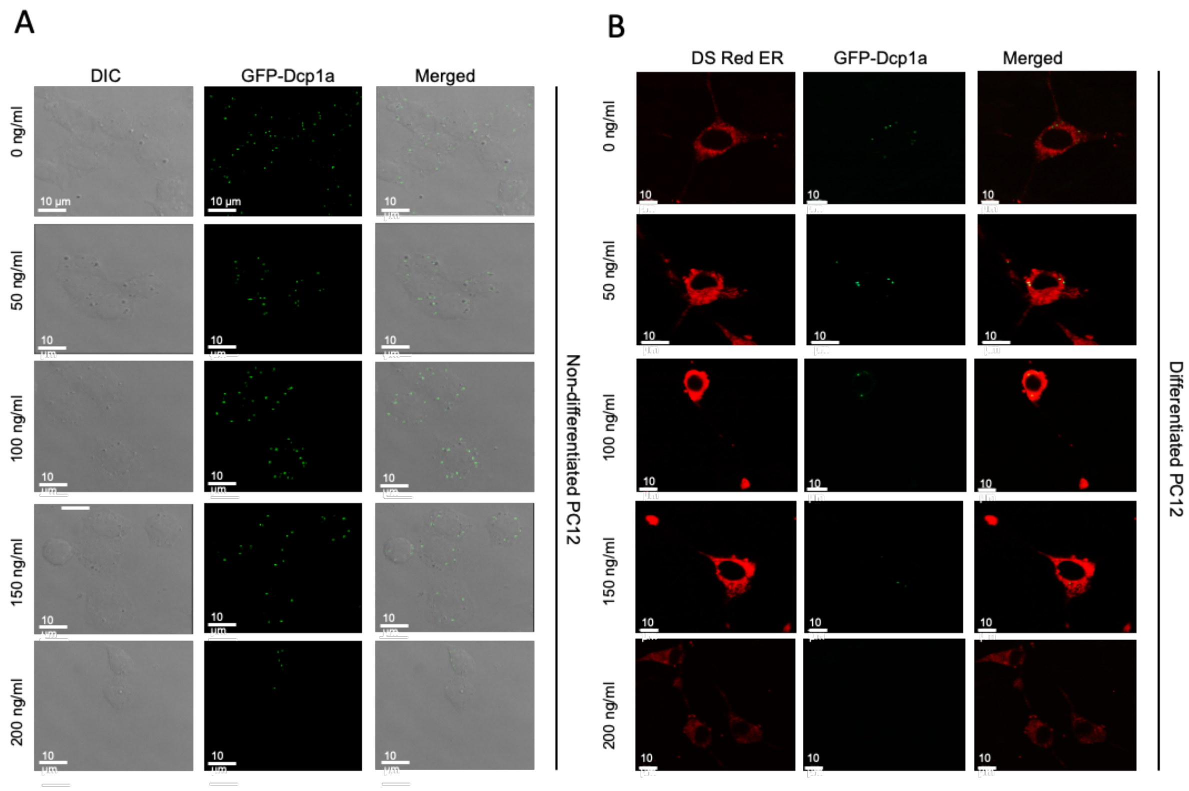


**Figure S1 Digitonin Permeabilization effects on internal P-bodies in PC12 cells. Related to Figure 1.**

**(A)** Confocal Images of GFP-Dcp1a (Green) expressing undifferentiated PC12 cells permeabilized with increasing concentration of Digitonin (0, 50, 100, 150, 200 ng/ml) for 10 minutes at 4 °C. Merged images are shown.

**(B)** Confocal Images of DS Red ER (Red) and GFP-Dcp1a (Green) expressing 72 hours differentiated (with 100 ng/ml NGF) PC12 cells permeabilized with increasing concentration of Digitonin (0, 50, 100, 150, 200 ng/ml) for 10 minutes at 4 °C. Merged images are shown.

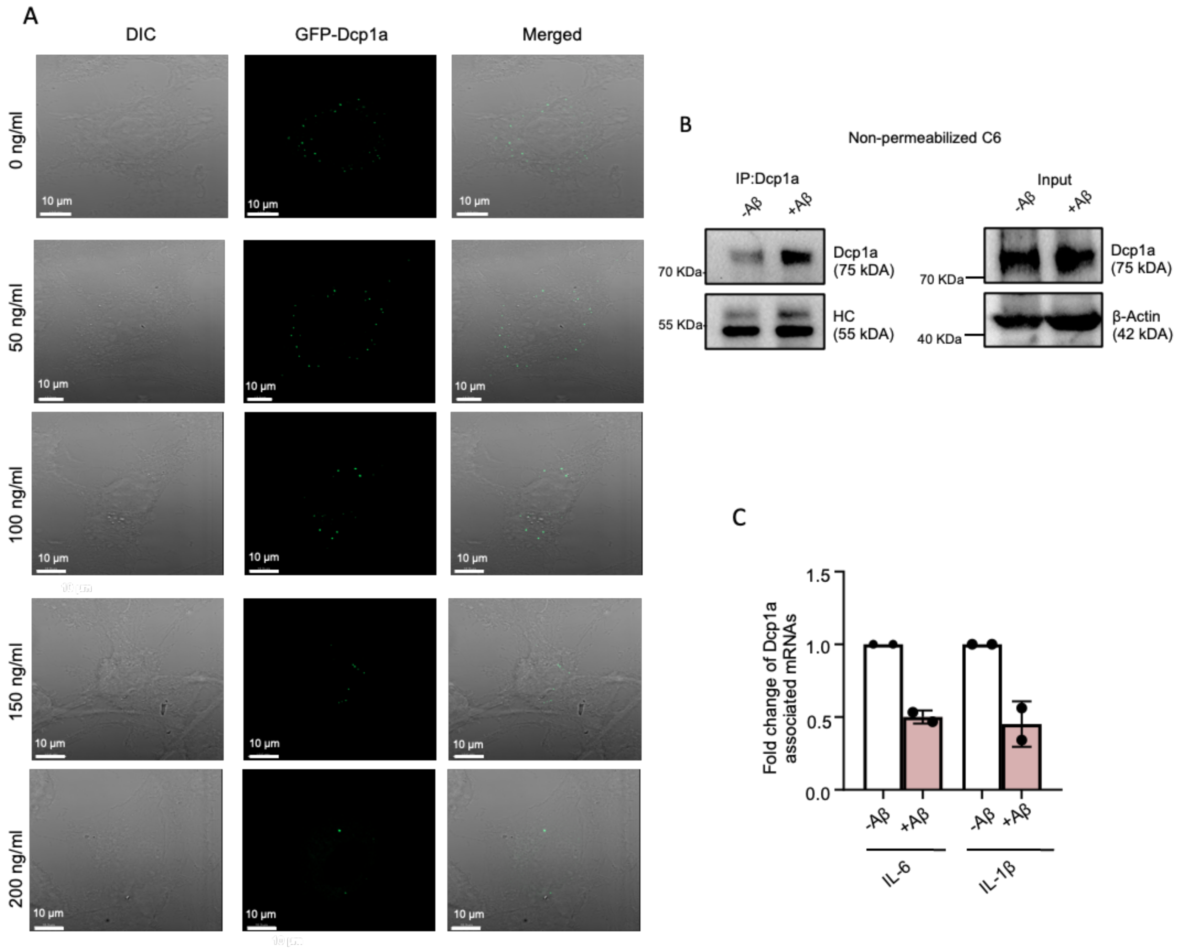


**Figure S2 Digitonin Permeabilization effects on internal P-bodies in PC12 cells. Related to Figure 1.**

**(A)** Confocal Images of GFP-Dcp1a (Green) expressing C6 cells permeabilized with increasing concentration of Digitonin (0, 50, 100, 150, 200 ng/ml) for 10 minutes at 4 °C. Merged images are shown.

**(B) (C)** Immunoprecipitation of total endogenous Dcp1a protein from both 0 or 2.5 µM Aβ $_{(1-42)}$ treated C6 cells, without permeabilization. Western blot images of pulled down Dcp1a level along with heavy chain and input Dcp1a level with β-Actin as loading control are shown. Molecular weight markers are also shown. (B). Followed by qRT-PCR based quantification of associated IL-6 and IL-1β mRNA levels normalized against relative pulled down Dcp1a levels (n=2 independent experiments) (C).

**Table S1: List of primary antibodies**

| Antigen name | Raised in | Source | Catalogue No. | Dilution for WB | Dilution for IF/IP |
|---|---|---|---|---|---|
| DDX6 | Rabbit | Bethyl | A300-461A | 1:10000 | - |
| DCP1A | Mouse | Abnova | H0055802-M06 | 1:1000 | 1:100 /1:100 |
| HA | Rat | Roche | 11867423001 | 1:1000 | 1:100 /1:100 |
| β-Actin (HRP) | Mouse | Sigma | A3854-200UL | 1:10000 | - |
| Myc | Mouse | Santa Cruz | (9E10):sc-40 | 1:1000 | - |
| β-Tubulin | Mouse | Sigma-Aldrich | T5201 | 1:6000 | - |
| GAPDH | Mouse | Sigma-Aldrich | G9295 | 1:6000 | - |
| eIF2C2 | Mouse | Abnova | H00027161-M01 | - | 1:100 |

**Table S2: List of secondary antibodies**

| Secondary Antibody | Source | Catalogue No. | Dilution for WB | Dilution for IF |
|---|---|---|---|---|
| HRP Goat anti-rat | Invitrogen | 62-9520 | 1:8000 | - |
| HRP Goat anti-rabbit | Invitrogen | #65-6120 | 1:8000 | - |
| HRP Goat anti-mouse | Invitrogen | #62-6520 | 1:8000 | - |
| Alexa Fluor 568 (mouse) | Invitrogen | A11004 | - | 1:500 |

**Table S3: List of plasmids and synthetic miRNA**

| Name | Source/Reference | Description |
|---|---|---|
| FLAG-HA-Ago2 | Kind gift from Tom Tuschl | Plasmid expressing FLAG and HA tagged human Ago2 |
| GFP-Dcp1a | Kind gift from Witold Filipowicz | Dcp1a cloned in frame with GFP |
| pCI-neo | Promega | pCI-neo expressing vector |
| Rheb-Myc | Kind gift from J. M. Backer | Myc tagged human Rheb expression plasmid |
| pDsRed2-ER | From Clonetech | ER targeting variant of DsRed |
| RL-IL-6 | Prepared by cloning | 3’UTR of IL-6 downstream of RL coding region in pCI-Neo vector |
| RL-IL-1β | Prepared by cloning | 3’UTR of IL-1β downstream of RL coding region in pCI-Neo vector |

**Table S4: List of mRNA real-time primers**

| Name | Primer | Sequence |
|---|---|---|
| GAP43 | Forward | 5’ ACGAGAAGAAGGGTGATGCA 3’ |
| | Reverse | 5’ CTTCGCCCTTCTTCTCCTCA 3’ |
| Neurofilament-M | Forward | 5’ AACGTCAAGATGGCTCTGGA 3’ |
| | Reverse | 5’ GTGTTGGACCTTGAGCTTGG 3’ |
| HuD | Forward | 5’ TCACCATTGACGGGATGACA 3’ |
| | Reverse | 5’ ACCTTGACGTTGTTCACTGC 3’ |
| HuR | Forward | 5’ TCAACTCCAGGGTCCTTGTG 3’ |
| | Reverse | 5’ GTTCTGGTTGGGATTGGCTG 3’ |
| GAPDH | Forward | 5’ CAGGGGGGAGCCAAAAGGG 3’ |
| | Reverse | 5’ CTTGGCCAGGGGTGCTAAGC 3’ |
| IL-6 | Forward | 5’ TACCCCAACTTCCAATGCTC 3’ |
| | Reverse | 5’ ACCACAGTGAGGAATGTCCA 3’ |
| IL-1β | Forward | 5’ GTGGATCCCAAACAATACCC 3’ |
| | Reverse | 5’ AACTATGTCCCGACCATTGC 3’ |
